# Patient-specific detection of cancer genes reveals recurrently perturbed processes in esophageal adenocarcinoma

**DOI:** 10.1101/321612

**Authors:** Thanos P. Mourikis, Lorena Benedetti, Elizabeth Foxall, Damjan Temelkovski, Joel Nulsen, Juliane Perner, Matteo Cereda, Jesper Lagergren, Michael Howell, Christopher Yau, Rebecca C. Fitzgerald, Paola Scaffidi, Francesca D. Ciccarelli, on behalf of the Oesophageal Cancer Clinical and Molecular Stratification (OCCAMS) Consortium

## Abstract

The identification of somatic alterations with a cancer promoting role is challenging in highly unstable and heterogeneous cancers, such as esophageal adenocarcinoma (EAC). Here we developed a machine learning algorithm to identify cancer genes in individual patients considering all types of damaging alterations simultaneously (mutations, copy number alterations and structural rearrangements). Analysing 261 EACs from the OCCAMS Consortium, we discovered a large number of novel cancer genes that, together with well-known drivers, help promote cancer. Validation using 107 additional EACs confirmed the robustness of the approach. Unlike known drivers whose alterations recur across patients, the large majority of the newly discovered cancer genes are rare or patient-specific. Despite this, they converge towards perturbing cancer-related processes, including intracellular signalling, cell cycle regulation, proteasome activity and Toll-like receptor signalling. Recurrence of process perturbation, rather than individual genes, divides EACs into six clusters that differ in their molecular and clinical features and suggest patient stratifications for personalised treatments. By experimentally mimicking or reverting alterations of predicted cancer genes, we validated their contribution to cancer progression and revealed EAC acquired dependencies, thus demonstrating their potential as therapeutic targets.

## INTRODUCTION

Genome instability enables the onset of several hallmarks of cancer as some of the acquired alterations confer selective advantages to the mutated cells, thus driving their outgrowth and eventual dominance (1). The identification of driver genes (the genes acquiring driver alterations) is therefore critical to fully understand the molecular determinants of cancer and to inform the development of precision oncology. Since driver genes are under positive selection during cancer progression, a reasonable assumption is that their mutation is observed more frequently than expected. Over the past years, large-scale cancer genomic studies have provided the required power to detect driver events recurring across samples with good statistical confidence (2,3). However, the full characterisation of driver events is particularly challenging when the genomic landscape of the cancer is highly variable and recurrent events are relatively rare.

One such cancer is esophageal adenocarcinoma (EAC), whose incidence in recent years has risen substantially in the western world (4). EAC exhibits high mutational and chromosomal instability leading to widespread genetic heterogeneity. In over 400 EACs sequenced so far, mutations in *TP53*, *CDKN2A*, *SMARCA4*, *ARID1A, SMAD4*, *ERBB2, MYD88, PIK3CA, KAT6A, ARID2* as well as amplifications of *VEGFA, ERBB2, EGFR, GATA4/6, CCNE1* are the most recurrent driver events (5-11). However, a significant fraction of patients is still left without known genetic determinants and often the number of identified drivers per sample is too low to fully explain the disease. Consequently, the molecular mechanisms that drive EAC have been difficult to characterise in full. This has a profound impact on the way in which EAC is currently diagnosed and treated. For example, phase III clinical trials with various targeted agents have failed to show benefits or reached inconclusive results (12-14).

Here we hypothesise that, alongside the critical role of recurrent and well-known drivers, complementary somatic alterations of other genes help cancer progression in individual patients. Therefore, the comprehensive characterisation of the full compendium of cancer drivers requires that both recurrent and rare events are considered. While recurrent drivers can be identified based on the frequency of their alterations, rare genes altered in few or even single patients are difficult to identify. To this aim, we developed sysSVM, an algorithm based on supervised machine learning that predicts cancer genes in individual patients. The rationale of sysSVM is that somatic alterations sustaining cancer affect genes with specific properties (15). It therefore uses these properties, rather than recurrence, to identify cancer genes.

We applied sysSVM to 261 EACs from the UK OCCAMS Consortium, which is part of the International Cancer Genome Consortium (ICGC). We first trained the classifier using 34 features derived from properties specific to known cancer genes and then prioritised 952 genes that, together with the known drivers, help promote cancer development across the whole EAC cohort. The large majority of these newly predicted ‘helper’ genes are rare or patient-specific. When analysing the biological pathways that they are involved in, helpers converge towards the perturbation of cancer-related processes such as intracellular signalling, cell cycle regulation, proteasome activity and Toll-like receptor signalling. We used the recurrence of process perturbation, rather than genes, to stratify the 261 EACs into six clusters that show distinct molecular and clinical features and suggest differential response to targeted treatment.

## RESULTS

### The landscape of recurrent and rare EAC genes

sysSVM applies machine learning to predict altered genes contributing to cancer in individual patients based on the similarity of their molecular and systems-level properties to those of known cancer genes (Supplementary text). Molecular properties include somatic alterations with a predicted damaging effect on the protein function (gene gains and losses, translocations, inversions, insertions, truncating and non-truncating damaging alterations and gain of function mutations) as well as the overall mutation burden and the gene copy number (Supplementary Table 1). Systems-level properties are genomic, epigenomic, evolutionary, network and gene expression features that distinguish cancer genes from other genes. They include gene length and protein domain organisation (15,16), gene duplicability (17,18), chromatin state (19), connections and position of the encoded proteins in the protein-protein interaction network (17), number of associated regulatory miRNAs (18), gene evolutionary origin (18) and breadth of gene expression in human tissues (15,16) (Supplementary Table 1).

sysSVM is composed of three steps (Figure 1A, Supplementary text). In step 1, 34 features that describe the gene molecular and systems-level properties are mapped to all altered genes in each patient. In step 2, known cancer genes altered in the patient cohort are used to run a set of three-fold cross validations and identify the best models in four kernels (linear, sigmoid, radial, polynomial). In step 3, these best models used for training and prediction. All altered genes except known cancer genes are first scored in each patient individually by combining the predictions of the four kernels and then ranked according to the resulting score. Since the hypothesis is that the strength of the contribution of a gene to cancer depends on how similar its properties are to those of known cancer genes, the top scoring genes in each patient are the most likely contributors to cancer progression. The overall results are combined to obtain the final list of predicted cancer genes.

**Figure 1.**
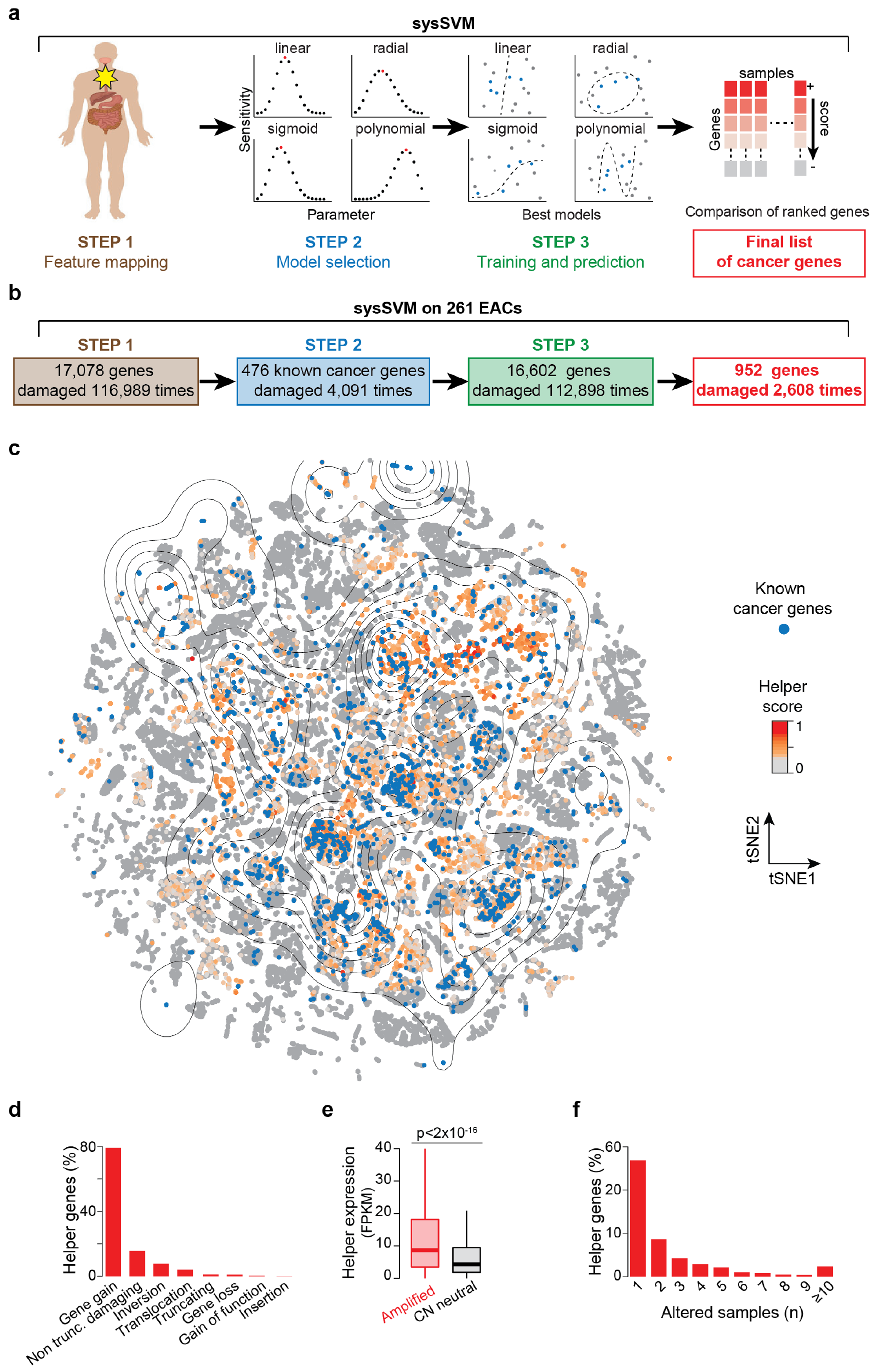
Cancer helper genes in 261 EACs. **a.** Schematic workflow of the sysSVM algorithm. **b.** Application of sysSVM to 261 EACs. Genes with somatic damaging alterations (n=116,989) were extracted from 261 EACs and divided into training (known cancer genes, blue) and prediction (rest of altered genes, green) sets. sysSVM was trained on the properties of known drivers and the best models were used for prediction. All altered genes were scored in each patient individually and the top 10 hits were considered as the cancer helper genes in that patient, for a total of 2,608 helper genes, corresponding to 952 unique hits (red). **c.** t-distributed Stochastic Neighbour Embedding (t-SNE) plot of 116,989 altered genes in 261 EACs. Starting from the 34 properties used in sysSVM, a 2-D map of the high-dimensional data was built using Rtsne package (https://github.com/jkrijthe/Rtsne) in R. Curves are coloured according to the density of 4,091 known cancer genes (blue) used as a training set and the rest of altered genes are coloured according to their sysSVM score. **d.** Distribution of damaging alterations in 952 cancer helpers. Overall, these genes acquire 2,608 damaging alterations. **e.** Expression of helper genes in EACs where they are amplified as compared to EACs where they are copy number neutral. FPKM values from RNA-Seq were available from 92 EACs. Out of the 952 helper genes, 389 had at least one amplification across these samples, for a total of 751 amplification events. Significance was assessed using the Wilcoxon rank-sum test. **f.** Recurrence of cancer helpers across 261 EACs. Only samples acquiring alterations with a damaging effect are considered.

We applied sysSVM to 261 EACs from OCCAMS which are part of the ICGC dataset (Figure 1B, Supplementary Table 2). In step 1, we extracted 17,078 genes with predicted damaging alterations (median of 382 damaged genes per patient) and mapped their 34 features. We verified that there is no pairwise correlation between these features (Supplementary Figure 1). Moreover, 476 known cancer genes (20) altered in the 261 EACs (Supplementary Table 3) tend to cluster in distinct regions of the feature space (Supplementary Figure 2). This confirms that these features distinguish cancer genes from other genes. In step 2, we ran 10,000 iterations of a three-fold cross validation using the 476 known cancer genes and combined the results to obtain 500 best models for each kernel (Supplementary Table 4, Methods). In step 3, we trained the four classifiers with these best models and used them to score and rank the remaining 16,602 altered genes in each patient. Since the gene score reflects a gradient between driver and passenger activity, we considered the top 10 scoring genes in each EAC as the main cancer contributors for that patient. We verified that the main findings of our study hold true if we apply higher or lower cut offs (see below). Overall, this produced 500 lists of top 10 scoring genes in each sample (Supplementary Table 4, Methods). We considered the list of 952 genes that occurred most frequently as the final set of predicted cancer genes (Supplementary Table 5). Since our hypothesis is that these genes help the known drivers to promote cancer, we define them as ‘helper genes’.

Helper genes localise to the same high-density regions of known cancer genes (Figure 1C), with lower scoring genes being further away from these regions (Supplementary Figure 3>). This indicates that the properties of top scoring genes indeed resemble those of known cancer genes. Consistent with the prevalence of gene amplification in EAC (Supplementary Table 1), the vast majority of the 952 helpers undergo copy number gain (Figure 1D), resulting in their increased expression (Figure 1E). Despite the majority of helpers being rare or patient-specific (Figure 1F), a few of them are altered in more than 10% of EACs (Supplementary Table 5). These genes are usually associated with frequently occurring amplification events (10) (Supplementary Figure 4).

We assessed the robustness of our predictions in several ways. First, we evaluated the performance of sysSVM using two independent cohorts, namely 86 EACs from The Cancer Genome Atlas (TCGA) and 21 EACs from another study (21) (Supplementary Table 2). We scored all altered genes, including known cancer genes (20), in each of the 107 EACs independently, using the four best models trained on the ICGC cohort. In both datasets, known cancer genes have significantly higher scores than the rest of the altered genes (Supplementary Figure 3), indicating that sysSVM is able to recognise them as major cancer contributors. Second, we searched whether any of the 952 helpers were previously identified as cancer genes. We found that 41 of them (4%) have recently been added as known cancer genes to the Cancer Gene Census (22) and 171 helper genes (18%) have been predicted as candidate cancer genes in various cancers, including EAC (5) (Supplementary Table 5). Third, we searched for possible false positive predictions using two lists. The first was composed of 49 genes predicted as false positives of recurrence-based methods (5) and found only three helpers (*PCLO*, *CNTNAP2* and *NRXN3*) (Supplementary Table 5). Interestingly, *PCLO* has recently been shown to exert an oncogenic role in esophageal cancer by interfering with *EGFR* signalling (23). The second list was a manually curated set of 488 putative false positives (3) where we found 44 helpers (4.6% of the total). This is less than the fraction of known cancer genes (20) present in the same list of false positives (46/719, 6.4%). Altogether these analyses indicate that sysSVM robustly predicts cancer genes in multiple patient cohorts, with a minimal false positive rate.

### Helper genes converge to perturb related biological processes

In order to gather a comprehensive characterisation of the molecular determinants of EAC, we analysed the biological processes perturbed by helpers compared to drivers. We manually reviewed all 476 known cancer genes (20) with damaging alterations in the OCCAMS cohort and retained 202 of them based on the concordance between the type of acquired modification and the literature evidence of their cancer role (Methods, Supplementary Table 3). The median number of drivers per EAC is in accordance with recent estimates (24,25) and the majority of them undergo gene amplification (Supplementary Figure 4). We then performed two independent gene set enrichment analyses, one with the 202 known drivers and one with the 952 helpers, to dissect their relative functional contribution to EAC. This led to 212 and 189 enriched pathways out of the 1,877 tested, respectively (FDR <0.01, Supplementary Table 6, Supplementary Figure 4). Interestingly, the analysis of known drivers resulted in a higher number of enriched pathways than helpers, despite their lower number. This reflects the higher number of pathways that drivers map to (median of four pathways for known drivers and two pathways for helpers).

Seventy-three pathways (over 34%) enriched in known drivers are perturbed in more than 50% of EACs (Supplementary Table 6, Supplementary Figure 5). These ‘universal cancer pathways’ are involved in well-known cancer-related processes, such as intracellular signalling, cell cycle control, apoptosis and DNA repair, and are associated with the most recurrently altered known drivers (*TP53, CDKN2A, MYC, ERBB2, SMAD4, CDK6, KRAS*, Supplementary Table 3). Interestingly, 50 of the 73 (70%) are also enriched in helpers and 86 patients with altered helpers in a universal cancer pathway have no known drivers in that pathway (Figure 2A, Supplementary Table 6). This indicates that helpers often contribute to the perturbation of key cancer pathways and that their alteration may be sufficient for cancer development in the absence of known drivers.

**Figure 2.**
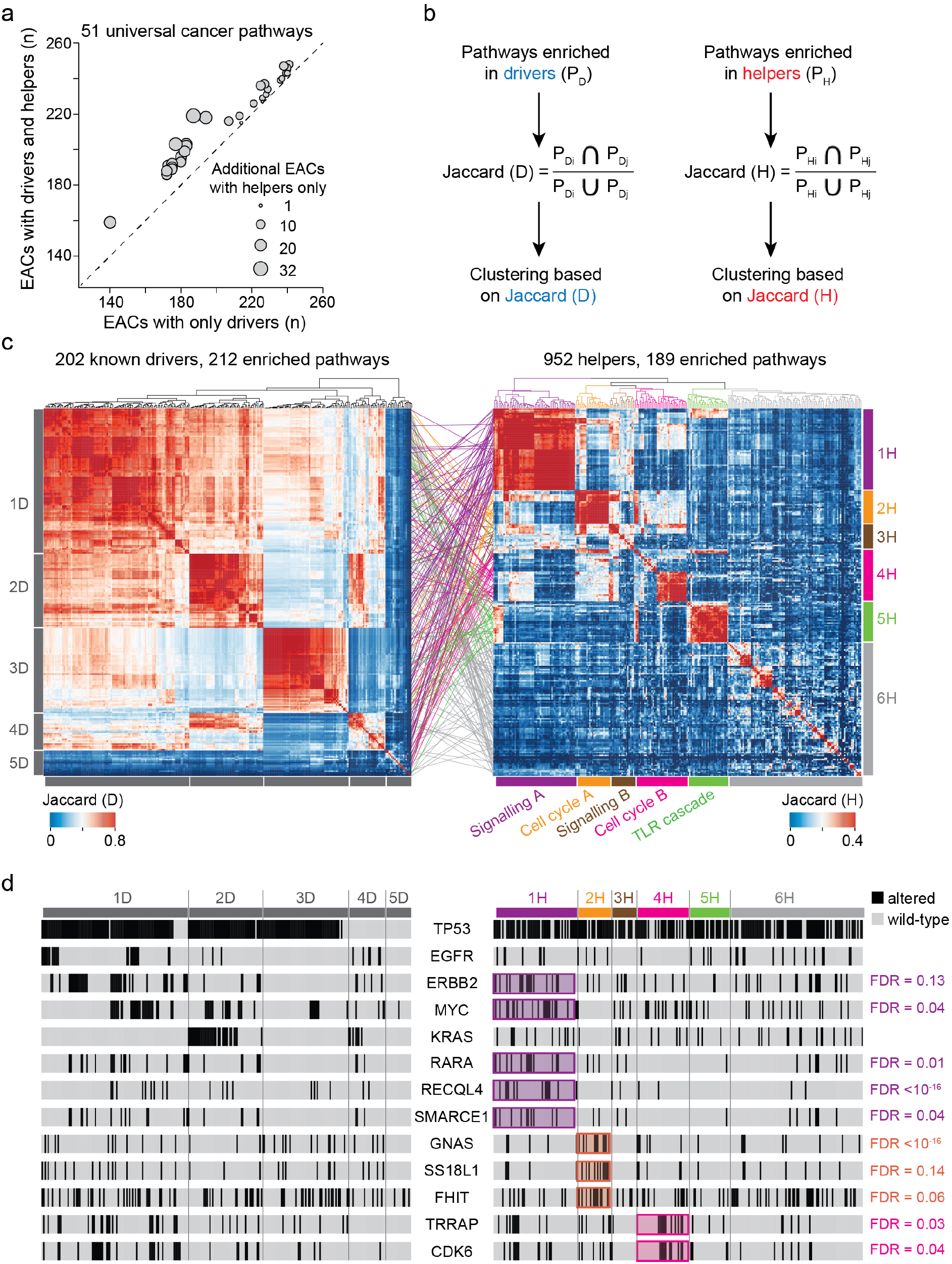
Perturbed processes in 261 EACs. **a.** Scatterplot of 51 universal pathways enriched in known drivers and helpers. For each pathway, the number of EACs with altered drivers and the number of EACs with altered drivers and helpers is shown. The size of dots is proportional to the additional EACs with perturbations in these pathways because of altered helpers only. **b.** Schematic of the procedure to cluster EACs according to pathways enriched in known drivers or helpers. Enriched pathways are mapped to individual EACs and the Jaccard index is calculated as the proportion of shared pathways over the total pathways in each pair of samples (*i, j*). Hierarchical clustering is then performed. **c.** Clustering of 261 EACs according to pathways enriched in known drivers and helpers. Five clusters were identified using known drivers (1D-5D) and six using helpers (1H-6H). Cluster-matching coloured lines show where EACs clustered by pathways enriched in helpers map in the driver clusters. **d.** Mutational status of selected known drivers across 261 EACs. Drivers enriched in clusters of helpers are highlighted. Significance was assessed using the Fisher’s exact test, after correcting for False Discovery Rate (FDR).

Next, we clustered EACs according to the proportion of perturbed pathways that they have in common (Methods, Figure 2B). When using pathways enriched in known drivers, we identified five well-supported clusters (1D-5D, Figure 2C, median silhouette score = 0.5, Supplementary Figure 6). These clusters are clearly driven by the mutational status of the most recurrent drivers. For example, *TP53* is altered in clusters 1D-3D, *EGFR*, *ERBB2* and *MYC* are altered in cluster 1D and *MYC* and *KRAS* are altered in cluster 2D (Figure 2 D, Supplementary Table 2). Samples in clusters 4D and 5D show an overall lower mutational burden (p = 0.03, Wilcoxon rank sum test), fewer known drivers and consequently a lower number of enriched pathways (p = 7×10^-6^, Wilcoxon rank sum test, Supplementary Figure 4).

When clustering EACs according to the pathways enriched in helpers, we identified six well-supported clusters (1H-6H, Figure 2C, median silhouette score = 0.3, Supplementary Figure 6). In this case, samples are brought together not by the recurrent alterations of known driver genes, but by several helpers mapping to the same or related pathways (Supplementary Table 2). For example, both clusters 1H and 3H show diffuse perturbations in intracellular signalling (Figure 2C), very often involving universal cancer pathways (Supplementary Table 6, Supplementary Figure 7). In more than 43% of EACs in both clusters, the perturbations in universal cancer pathways occur in patients with no drivers. Other pathways perturbed in cluster 1H, but not in 3H, involve cell cycle regulation, Toll-like receptor signalling and proteasome activity (Supplementary Table 2, Supplementary Figure 7). EACs in cluster 1H also have significant association with several known cancer drivers such as *RECQL4*, *RARA*, *MYC*, *SMARCE1* and *ERBB2* (Figure 2D), which are often but not always co-altered (Supplementary Figure 4). They have a prevalence of mutational signature S3 and are enriched in early (stage 2) tumours (Figure 3A). Patients in cluster 3H are instead enriched in tobacco smokers (Figure 3A).

**Figure 3.**
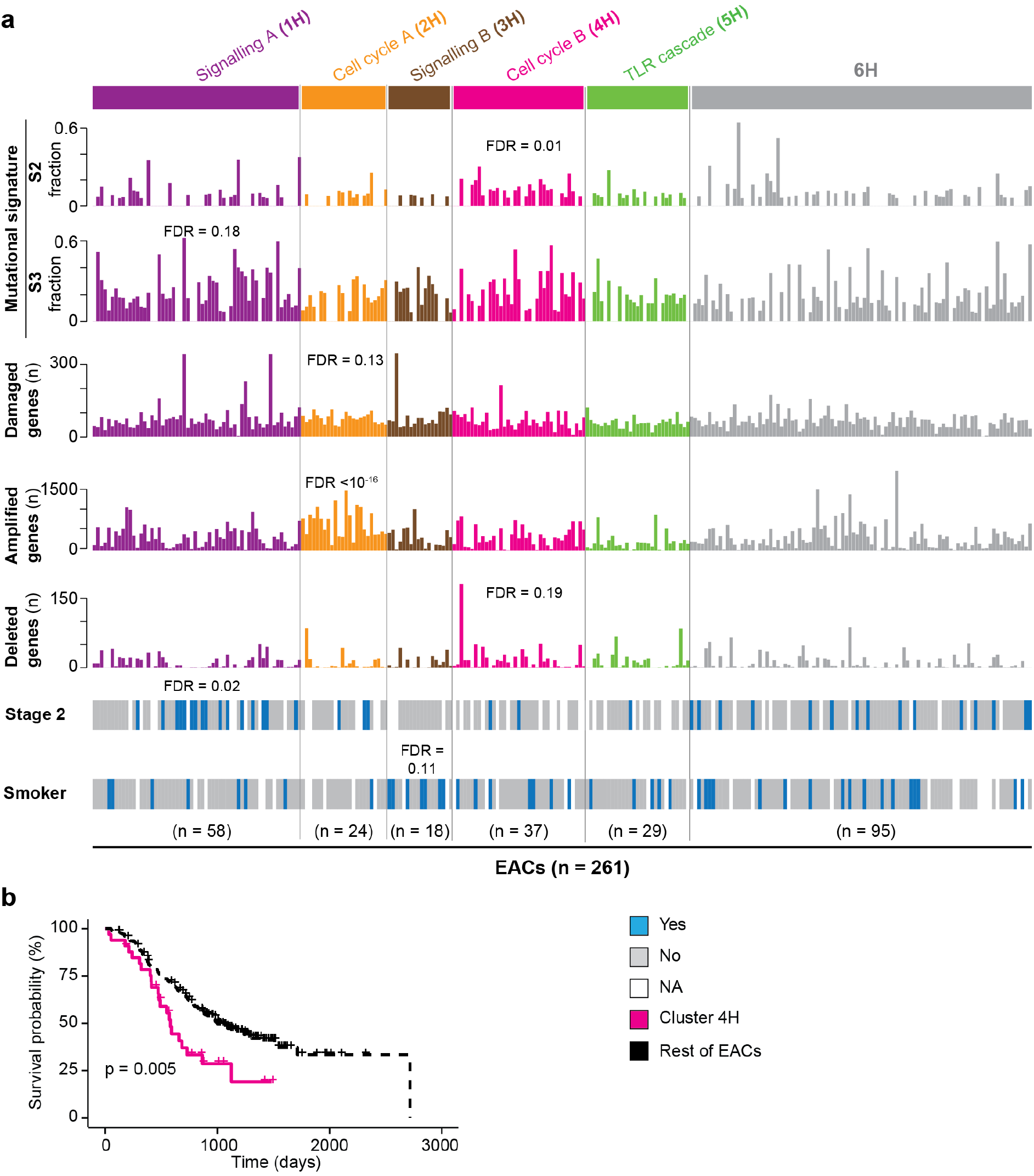
Features of EAC clusters driven by pathways enriched in helpers. **a.** For each helper cluster (1H-6H) indicated are the molecular features (mutational signatures, number of genes with damaging mutations, undergoing amplification or deletion), the distribution of stage 2 tumours and the tobacco smoking habits of the patients that show significant associations with one of the six clusters of helpers. Enrichment in number of altered genes, tumour staging and smoking habits was assessed using Fisher’s exact test. Distributions of mutational signatures were compared using Wilcoxon rank-sum test. FDR = false discovery rate after correction for multiple testing. **b.** Kaplan-Meier survival curves of EACs in cluster 4H (n = 37) and the rest of EACs (n = 224). Analysis was performed using survival and survminer R packages with default parameters. Significance was measured using the log-rank test

Similar to clusters 1H and 3H, the processes perturbed in clusters 2H and 4H are also functionally related, in this case to cell cycle regulation (Figure 2C). All EACs in cluster 2H have helpers involved in the regulation of G1/S transition (Supplementary Figure 7), such as members of the E2F family of transcription factors and their associated co-activators, competitors and downstream targets (Supplementary Table 2). Cluster 4H instead harbours perturbations in DNA replication, with alterations in the MCM complex, which is a downstream target of E2F (26,27). Dysregulation of E2F transcription factors or the MCM complex can induce increased genomic instability through either aberrant cell-cycle control or replicative stress (28,29). Consistently with this, EACs in clusters 2H and 4H are genomically unstable. Samples in 2H accumulate significantly more damaged and amplified genes, while those in 4H show significantly more deleted genes and are enriched in mutational signature 2 (Figure 3A). Cluster 2H also shows significant alterations of the known drivers *GNAS*, *SS18L1*, and *FHIT* (Figure 2D). *FHIT* is linked to increased genomic instability (30) and regulates the expression of cell cycle-related genes (31), therefore potentially affecting the G1/S transition pathways of this cluster. Cluster 4H shows frequent alterations in the known drivers *TRAPP* and *CDK6* (Figure 2D). The latter functions in various cell cycle-related pathways, including the mitotic G1/S phase pathway altered in 100% of cluster 4H (Supplementary Table 2, Supplementary Figure 7). Furthermore, patients in cluster 4H have a significantly lower survival compared to the rest of the cohort (Figure 3B). Interestingly, elevated expression of the MCM complex has been associated with poor patient survival in multiple tumour types (32). The perturbation of MCM proteins and their related pathways could therefore contribute to tumour aggressiveness and poor outcome among patients in this cluster. Finally, cluster 5H shows perturbations in the Toll-like receptor (TLR) signalling cascade (Supplementary Figure 7) that has recently been reported to be dysregulated in EAC (33).

Overall, clusters 1H to 5H account for 166 EACs (64% of the total cohort). The remaining 95 EACs in cluster 6H share fewer perturbed pathways, although 55 of them (58%) have alterations in Rho GTPase activity (Supplementary Figure 7) with frequent modifications of Rho GTPase effectors such as *ROCK1*, *PTK2*, *PAK1*, *LIMK1* and *NDE1* (Supplementary Table 2). EACs in the six clusters obtained using helpers are broadly dispersed in the clustering of known drivers (Figure 2C) indicating that helpers bring together patients with similar perturbed processes that cannot be appreciated when focussing only on recurrent drivers.

To test whether the clustering is affected by considering only the top 10 helper genes in each patient, we performed the same analysis considering as helpers the top five or top 15 scoring genes (528 and 1,297 unique genes, respectively). We found that the vast majority (99% and 77%) of the pathways enriched in these two datasets are also enriched when considering the top 10 helpers (Supplementary Figure 8). This indicates that the recurrently perturbed processes are highly overlapping. We then clustered EACs according to the proportion of common perturbed pathways and verified that the six clusters obtained using pathways enriched in top 10 genes recapitulated well the clusters obtained using pathways enriched in top five or 15 genes (Supplementary Figure 8). Therefore, the clustering is robust regardless of the applied ranking cut-off.

### Helper alterations contribute to cancer-related phenotypes and lead to dependence

To test the contribution of EAC helper genes towards the perturbation of cellular processes, we used two experimental approaches. In the first one, we assessed the consequences of altering representative helpers in FLO-1 cells. These are an EAC diploid cell line with no mutations or copy number alterations in any of the helpers selected for validation (34), thus allowing a clear evaluation of the effect of their alteration. We measured cell proliferation as a main hallmark of cancer (3,35) and also performed gene-specific assays. In the second approach, we assessed the effect of reverting the alteration of helpers on cell growth to evaluate the dependence of EAC on helper perturbations. In this case, we used EAC cell lines with alterations similar to those observed in patients.

We started by modifying the most commonly altered helpers in clusters 2H and 4H, namely *E2F1* (23 out of 24 samples in cluster 2H) and *MCM7* (18 out of 37 samples in cluster 4H, Supplementary Table 2). Both *E2F1* and *MCM7* are amplified in EACs (Supplementary Table 5) leading to significant gene overexpression (median two-fold increase, p = 6×10^-3^ and p=8×10^-3^, respectively Wilcoxon rank-sum test; Figure 4A). We therefore stably overexpressed *E2F1* and *MCM7* in FLO-1 cells to levels comparable to those observed in patients (Figure 4B). In both cases we observed significantly increased proliferation of overexpressing cells as compared to control cells (p=2×10^-4^ and p=9×10^-4^, respectively, two-tailed t-test; Figure 4C). Since E2F1 promotes cell cycle progression, we assessed DNA replication rate by measuring EdU incorporation during the cell cycle. We observed increased EdU intensity throughout S phase in *E2F1* overexpressing cells as compared to control cells (p<10^-4^, Mann Whiney U test; Figure 4D). This suggests that E2F1 may help cancer growth by promoting S phase entry. To assess the functional consequence of *MCM7* overexpression, we measured the loading of the MCM complex onto chromatin. We observed that *MCM7* overexpressing cells display a lower MCM fluorescence intensity overall as compared to control cells when staining the chromatin-bound fraction for either MCM7 or MCM3 (p<10^-4^, Mann-Whitney U test; Figure 4E, 4F). This suggests that less MCM complex is loaded onto chromatin by the start of S phase. Therefore, *MCM7* overexpression leads to both increased proliferation and perturbation of MCM complex activity. Finally, we reduced *MCM7* expression levels in MFD-1 cells that were recently derived from an EAC patient of our OCCAMS cohort (36). MFD-1 cells have four-fold higher *MCM7* basal expression compared to the diploid FLO-1 cells (Figure 4G). We therefore used doxycycline-inducible shRNA lentiviral vector (Supplementary Table 7) to reduce *MCM7* expression in MFD-1 cells to the level of FLO-1 cells (Figure 4H). This led to a significant decrease in cell proliferation (p = 2×10^-5^, two-tailed t-test; Figure 4I), indicating that MFD-1 cells rely on *MCM7* overexpression for their faster growth.

**Figure 4.**
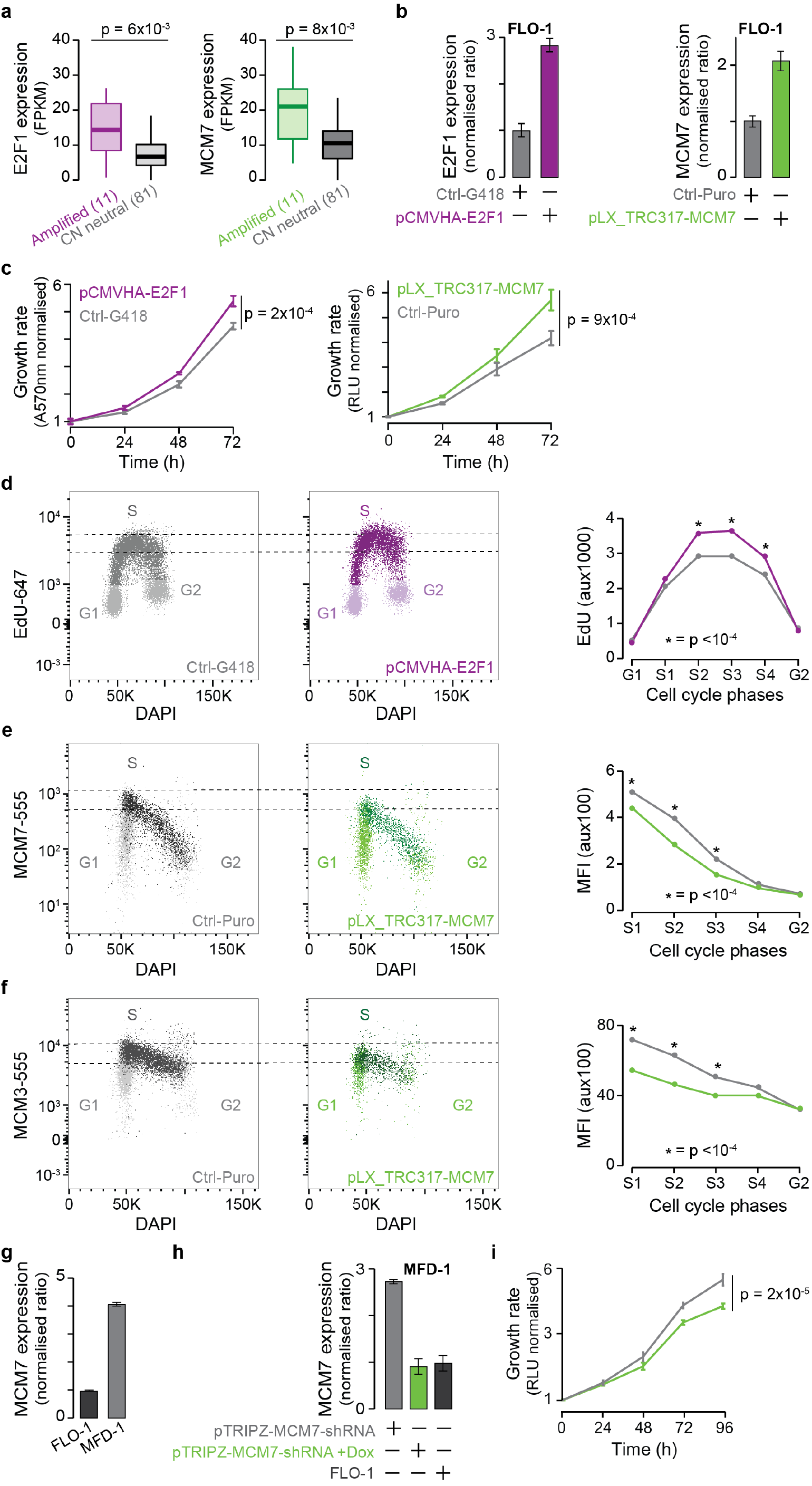
Cancer helper role of *E2F1* and *MCM7*. **a.** *E2F1* and *MCM7* expression in EACs where they are amplified (11 samples each) as compared to EACs where they are copy number neutral (81 samples each). Significance was assessed using the Wilcoxon rank-sum test. **b.** *E2F1* and *MCM7* mRNA expression in FLO-1 cells assessed by qRT-PCR. Expression was relativised to β-2-microglobulin and normalised to control cells. **c.** Proliferation curve of FLO-1 cells overexpressing *E2F1* or *MCM7* as compared to the corresponding control cells. **d.** Assessment of EdU (5-ethynyl-2’-deoxyuridine) incorporation by flow cytometry in *E2F1* overexpressing cells as compared to control cells. Cells were separated into G1, S and G2 phases, and S phase cells were subdivided into 4 gates from early to late S phase (S1-S4, Supplementary Figure 9). The geometric mean fluorescence intensity of EdU was measured for the cells in each gate and differences between EdU intensity were assessed using the Mann-Whitney U test. Three biological replicates were performed and a representative experiment is shown. Quantification of MCM complex loading onto chromatin in *MCM7* overexpressing or control cells via staining of MCM7 (**e**) or MCM3 (**f**). Cells were pulsed with EdU, and chromatin fractionation was performed before staining for MCM7 or MCM3 to detect the MCM complex bound to chromatin. Cells were separated into cell cycle phases using EdU and DAPI intensity (see Methods and Supplementary Figure 9). MCM7 or MCM3 fluorescence intensity during S phase illustrates the unloading of the MCM complex from chromatin. The geometric mean fluorescence intensity of MCM staining was measured for the cells in each cell cycle gate and differences in MCM intensity were assessed using Mann-Whitney U test. Representative data from one of three biological replicates are shown. Pseudocolour plots corresponding to panels E, F and G are shown in Supplementary Figure 9. **g.** *MCM7* mRNA expression levels in MFD-1 and FLO-1 cells. Expression was relativised to β-2-microglobulin and normalised to FLO-1 cells. **h.** *MCM7* expression levels in MFD-1 cells after transduction with a lentiviral vector carrying an inducible shRNA against *MCM7*. Expression was assessed in the absence of doxycycline and after 96 hours of doxycycline treatment, relativised to β-2-microglobulin and normalised to FLO-1 cells. **i.** Proliferation curve of MFD-1 cells with or without doxycycline-induced *MCM7* knockdown. For all qRT-PCR experiments, two biological replicates were performed, with reactions performed in triplicate. For all proliferation assays, at least two biological replicates were performed, each with four technical replicates. Proliferation was assessed every 24 hours and each time point was normalised to time zero. Mean values at 72 hours were compared by two-tailed Student’s t-test.

Next, we evaluated the role of rare helpers that were altered in a low fraction of patients. First, we tested *NCOR2* that is altered in eight EACs across five of the six clusters (Supplementary Table 5). *NCOR2* is part of the nuclear receptor corepressor complex that favours global chromatin deacetylation and transcriptional repression (37,38) (Figure 5A). Consistently with the suggested tumour suppressor role of *NCOR2* in lymphoma and prostate cancer (39,40), the most frequent *NCOR2* alterations in EAC lead to a loss of function. To reproduce these alterations, we edited the gene in FLO-1 cells using a vector-free CRISPR system (41). Three pooled crRNAs were co-transfected with Cas9 and the tracrRNA (Methods, Supplementary Table 7) and the editing was confirmed and quantified using Miseq (Figure 5B). We observed a 1.3-fold increase in proliferation in the edited cells compared to the control cells (p = 3×10^-3^, two-tailed t-test test; Figure 5C).

**Figure 5.**
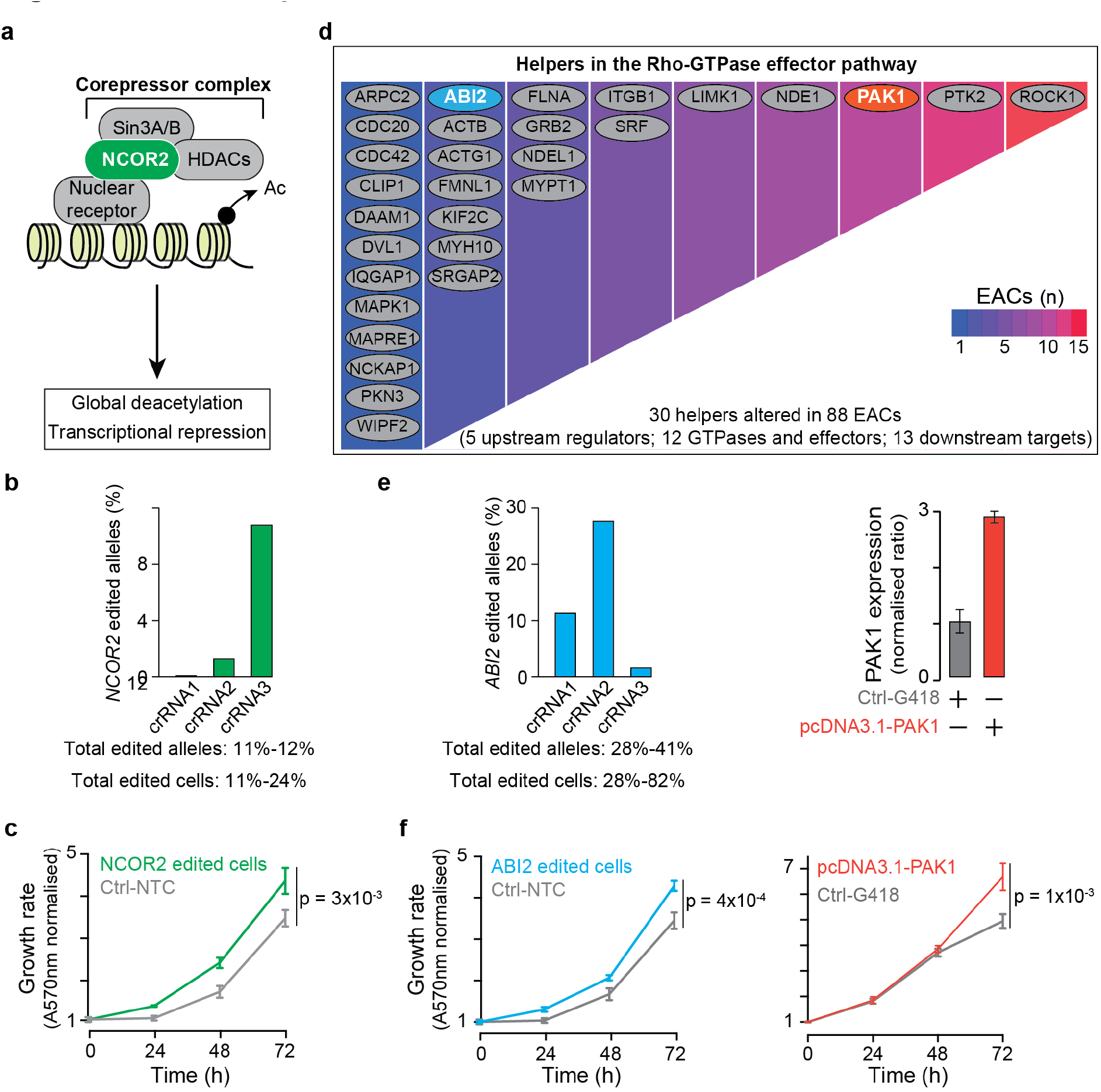
Cancer helper role of *NCOR2*, *ABI2*, and *PAK1*. **a.** Function of NCOR2 as part of the nuclear receptor co-repressor complex, whose activity results in chromatin deacetylation and transcriptional repression. **b.** Editing of the *NCOR2* gene using three pooled crRNAs where cells are transiently co-transfected with Cas9 protein, crRNAs and tracrRNA (41). The editing efficiency was measured using Miseq and the range of edited alleles and cells was derived considering the two opposite scenarios where all three crRNAs edit the same alleles/cells or different alleles/cells, respectively. **c.** Proliferation curve of *NCOR2* or NTC edited FLO-1 cells. Proliferation was assessed every 24 hours and each time point was normalized to time zero. Mean values at 72 hours were compared by two-tailed Student’s t-test. Three biological replicates were performed, each with four technical replicates. **d.** Manual curation of the helpers contributing to the Rho-GTPase effectors pathway. Heatmap indicates the number of samples with alterations in each gene. ABI2 (blue) and *PAK1* (red) were selected for experimental validation. **e.** Induced alterations in *ABI2* and *PAK1* genes. Editing of *ABI2* was performed and assessed as described for *NCOR2*. *PAK1* mRNA expression in FLO-1 cells was assessed by qRT-PCR, relativised to β-2-microglobulin and normalised to control cells. Experiments were done in triplicate in two biological replicates. **f.** Proliferation curves of FLO-1 cells after *ABI2* editing or *PAK1* overexpression. Three biological replicates were performed, each with four technical replicates. Proliferation was assessed every 24 hours and each time point was normalised to time zero. Mean values at 72 hours were compared by two-tailed Student’s t-test.

Then, we tested the effect of altering members of the Rho GTPase effector pathway, which is pervasively perturbed in all six clusters, often through patient-specific alterations (Figure 5D, Supplementary Table 2). As representatives of this pathway we modified *ABI2* and *PAK1*, which undergo damaging alterations and amplification in one and nine EACs, respectively (Supplementary Table 5). We therefore edited *ABI2* and overexpressed *PAK1* as described above (Supplementary Table 7, Figure 5E). In both cases we observed significantly increased proliferation as compared to control cells (ABI2: p = 4×10^-4^, PAK1: p = 1×10^-3^ two-tailed t-test; Figure 5F).

Finally, we focussed on *PSMD3* that encodes a subunit of the regulatory 19S proteasome complex. *PSMD3* is amplified and overexpressed in three EACs of cluster 1H, which overall contains 14 samples with alterations in six proteasome subunits (Figure 6A and Supplementary Table 5). We identified three EAC cell lines (MFD-1, OE19 and OE33) showing higher basal expression of *PSMD3* compared to FLO-1 (2-, 3- and 4-fold increase respectively, Figure 6B). Using a doxycycline-inducible lentiviral shRNA vector (Supplementary Table 7s), we reduced *PSMD3* expression in MFD-1, OE19 and OE33 cells to levels equivalent to those of FLO-1 (Figure 6C). In all three cell lines we observed a significant reduction in cell proliferation following the reduction of *PSMD3* expression (MFD-1: p = 4×10^-8^; OE19: p = 2×10^-8^; OE33: p = 6×10^-3^, two-tailed t-test; Figure 6D). The effect was particularly strong in OE19, where the reduction of *PSMD3* expression to diploid levels arrested cell growth completely. In MFD-1 and OE33 it led to 1.3- and 1.2-fold reduction of cell growth (Figure 6D). This suggests that the extent of EAC reliance upon helper alterations is at least partially context dependent.

**Figure 6.**
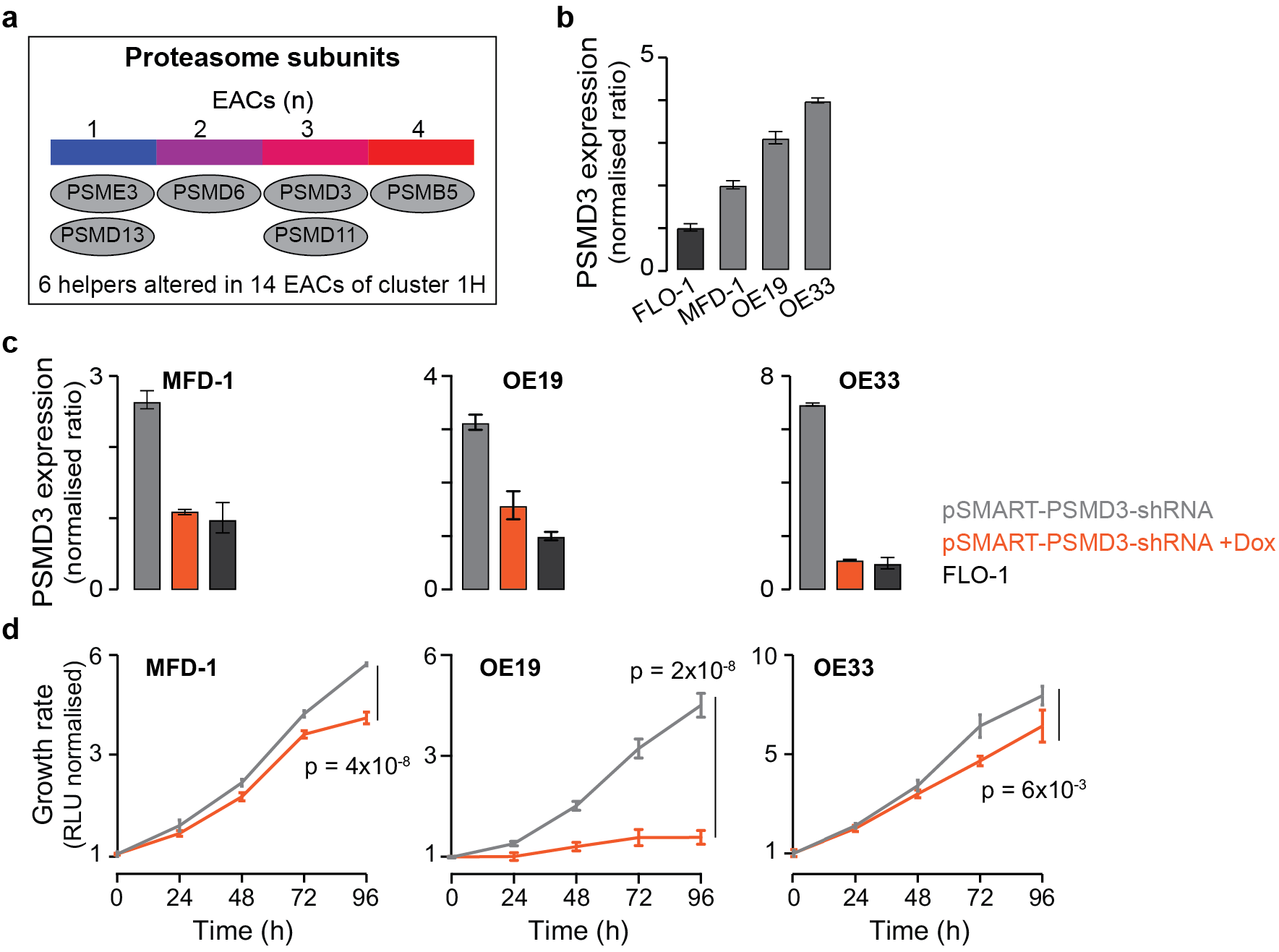
EAC cell dependence on *PSMD3* alteration. **a.** Heatmap of proteasome subunits predicted as helpers in 261 EACs. **b.** *PSMD3* basal mRNA expression levels in FLO-1, MFD-1, OE19 and OE33 cells. Expression was relativised to β-2-microglobulin and normalised to FLO-1 cells. **c.** *PSMD3* expression levels in MFD-1, OE19 and OE33 after transduction with a lentiviral vector carrying an inducible shRNA against *PSMD3*. Expression was assessed in absence of doxycycline and after 96 hours of doxycycline treatment, relativised to β-2-microglobulin and normalised to FLO-1 cells. **d.** Proliferation curves of MFD-1, OE19 and OE33 cells with or without doxycycline treatment to reduce *PSMD3* expression to levels comparable to those of FLO-1 cells. For all proliferation assays, at least two biological replicates were performed, each with four technical replicates. Proliferation was assessed every 24 hours and each time point was normalised to time zero. Mean values at 96 hours were compared by two-tailed Student’s t-test.

Taken together, our experimental data indicate that, independently of the alteration frequency, the modification of helpers positively affects EAC cell growth. Moreover, we provide evidence that EAC cells become addicted to helper alterations, suggesting that targeting helpers, or the pathways in which they act, could reduce EAC progression.

## DISCUSSION

Most state-of-the-art approaches to discovering cancer driver events rely on the detection of positively selected alterations of genes that are functionally beneficial for cancer development (3,24). Even ratiometric methods based on gene properties (42) ultimately assess the effect of positive selection and distinguish the few selected drivers from the many passenger events. As a result, the discovery of cancer drivers is biased towards genes that are frequently altered across patients. This poses significant limitations for cancers such as EAC that have a highly variable but mostly flat (*i.e.* with few recurrent events) mutational landscape. In support of this, the overall selection acting on esophageal cancer genomes is among the lowest across cancer types (24), despite a median of 382 damaged genes per EAC (Supplementary Table 2). This indicates that the exclusive focus on genes under strong selection is likely to return only a partial representation of the genes involved in EAC.

To overcome these limitations, our machine learning approach sysSVM ranks somatically altered genes that are relevant to cancer development based on their properties rather than mutation recurrence. Another advantage is that sysSVM considers all types of gene alterations (SNVs, indels, CNVs, and structural variations) simultaneously. Therefore, it provides a comprehensive overview of the genetic modifications that play a cancer promoting role in individual patients. When applied to 261 EACs, sysSVM prioritises 952 altered genes that, together with known drivers, help cancer progression. This large number of helper genes is in agreement with the recent observation of a positive correlation between mutational burden and number of driver genes, which is only partially explained by a sample size effect (3). We speculate that this positive correlation may indicate that the number of functionally relevant genes increases with the number of altered genes.

The heterogeneous landscape of EAC cancer genes is substantially reduced by considering the perturbed biological processes they act on (Figure 2C). Most of these processes are well-known contributors to cancer development, including intracellular signalling, cell cycle control, and DNA repair (Supplementary Table 6). Interestingly, while the known drivers tend to encode upstream players in these pathways, helpers are often downstream effectors. For example, we found several Rho GTPase effectors (Figure 5D, Supplementary Table 5) or genes downstream of previously reported EAC drivers in the Toll-like receptor cascade (SupplementaryTable 5). This supports a more local role of helpers in contributing to cancer at the single patient level, possibly by sustaining or complementing the function of drivers. In this respect, helpers are conceptually similar to mini-drivers (43).

Analysing the pathways disrupted by helpers allows the division of the 261 EACs into six functional clusters that often are closely related in function. For example, two of these clusters (1H and 3H) share perturbations in intracellular signalling. Similarly, clusters 2H and 4H show perturbations of processes involved in cell cycle, namely S-phase entry and DNA replication. Consistent with this, they bring together the most genomically unstable samples. By experimentally mimicking the amplification of *E2F1* (representative of cluster 2H) and *MCM7* (representative of cluster 4H), we induced increased proliferation in EAC cells (Figure 4C). We also provide evidence that E2F1 increases proliferation by promoting S phase entry (Figure 4D). Interestingly, *MCM7* overexpression resulted in a reduction of MCM complex loading onto chromatin (Figure 4E and 4F), maybe due to a stoichiometric imbalance of complex subunits. This may indicate that MCM7 promotes cell growth through a separate mechanism besides its function in the MCM complex. For example, MCM7 interacts with the tumour suppressor protein Rb, a well-characterised inhibitor of E2F1 (44). It is possible that *MCM7* overexpression may sequester Rb away from E2F1, thereby promoting E2F1-mediated cell cycle progression. Moreover, reducing *MCM7* expression levels in cells with high basal expression led to decreased cell proliferation, showing not only the contribution of *MCM7* alteration to EAC growth but also the dependence of cancer cells on its high expression.

We also confirmed the cancer promoting role of very rare helpers, such as *ABI2*, *NCOR2* and *PAK1* that are altered between 1% and 4% of EACs (Figure 5C and 5F). Therefore, irrespective of the frequency, helpers have a substantial impact on the progression of the cancer where their alteration occurs. This may indicate new, patient-specific gene dependencies and suggest possible stratifications that could inform the selection of targeted treatments. For example, 14 samples of cluster 1H have alterations of several proteasome subunits (Figure 6A, Supplementary Table 2). Experimentally reverting the expression of the proteasome subunit PSMD3 to diploid levels resulted in reduced cell growth in three different EAC cell lines (Figure 6D). This indicates that EACs become addicted to helper alterations and vulnerable to its inhibitions. Interestingly, proteasome inhibition has been shown to have a synergic effect in combination with ERBB2 inhibitors (45). Since *ERBB2* is also significantly altered in cluster 1H (Figure 2D), a combined therapy may be beneficial to patients in this cluster.

In summary, we provide one of the first attempts to extend the discovery of acquired perturbations contributing to cancer beyond those of recurrent drivers. Additional efforts are required to fully exploit the potential of these approaches to offer a more comprehensive view of the molecular mechanisms behind cancer and to guide novel clinical interventions.

## METHOD

### Annotation of molecular properties

Data on somatic single nucleotide variations (SNVs), small insertions and deletions (indels), copy number variations (CNVs), structural variations (SVs), and mutational signatures for 261 EACs were obtained from ICGC and analysed as previously described (10) (Supplementary Table 2). Briefly, SNVs and indels were called using Strelka v.1.0.13 (46) and subsequently filtered as previously described (10). For CNVs, the absolute copy number for each genomic region was obtained from ASCAT-NGS v.2.1 (47) after correction for tumour content, using read counts at germline heterozygous positions as derived from GATK v.3.2-2 (48). To account for the high number of amplifications occurring in EAC, copy number gains were corrected by the ploidy of each sample as estimated by ASCAT-NGS. A gene was assigned with the copy number of a CNV region if at least 25% of its length was contained in that region. SVs (gene translocations, inversions, insertions) were identified from discordant read pairs using Manta (49) after excluding SVs that were also present in more than two normal samples of a panel of 15 esophagus and 50 blood samples (10). In the case of TCGA validation cohort, SNVs, indels, and CNVs were derived from level 3 TCGA annotation data of 86 EACs (https://portal.gdc.cancer.gov/projects/TCGA-ESCA, Supplementary Table 2). In the case of 21 EACs from a previous study (21), SNVs, indels, and CNVs were called as described for the ICGC samples (Supplementary Table 2). The distribution of variant allele frequency of SNVs and indels across all samples was used to remove outliers likely indicating sequencing or calling artefacts. Variants with <10% frequency and indels longer than five base pairs were also removed. For CNVs, genomic regions were considered as amplified or deleted if their segment mean was higher than 0.3 or lower than −0.3, respectively, capping the segment mean to 1.5 to avoid hypersegmentation (50). A gene was considered as amplified or deleted if at least 25% of its length was contained in a CNV region and the resulting copy number (CN) was estimated as:

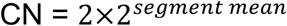

No SV data were available for the validation cohorts.

Since only genes with predicted damaging alterations were used as input for sysSVM, further annotation for the variant damaging effect was performed. Stopgain, stoploss, frameshift, nonframeshift, nonsynonymous, and splicing SNVs and indels were annotated using ANNOVAR (December 2015) (51). All truncating alterations (stopgain, stoploss, and frameshift mutations) were considered as damaging. Nonframeshift and nonsynonymous mutations were considered as non-truncating damaging alterations if predicted by at least five of seven function-based methods (SIFT (52), PolyPhen-2 HDIV (53), PolyPhen-2 HVAR (53), MutationTaster (54), MutationAssessor (55), LRT (56) and FATHMM (57)) or by two out of three conservation-based methods (PhyloP (58), GERP++ RS (59), SiPhy (60)), using the scores from dbNSFP v.3.0 (61). Splicing modifications were considered as damaging if predicted by at least one of the two ensemble algorithms as implemented in dbNSFP v3.0. Putative gain of function alterations were predicted with OncodriveClust (62) with default parameters and applying a false discovery rate of 10%. The transcript lengths to estimate mutation clustering were derived from the refGene table of UCSC Table Browser (https://genome.ucsc.edu/cgi-bin/hgTables). Gene gains, homozygous losses, translocations, inversions, insertions were always considered as putative damaging alterations.

Overall, 17,078 genes had at least one damaging alteration, for a total of 116,989 redundant damaged genes across 261 EACs (Supplementary Table 1). Of these, 476 were known cancer genes (20), corresponding to 4,091 redundant genes (Supplementary Table 2). For all 17,078 genes, the total number of exonic alterations (silent and nonsilent) and the somatic copy number were used as additional molecular features in sysSVM.

### Annotation of systems-level properties

Protein sequences from RefSeq v.63 (63) were aligned to the human reference genome assembly GRCh37 to define unique gene loci as previously described (16). The length of the longest coding sequence was taken as the gene length. Genes aligning to more than one gene locus for at least 60% of the protein length were considered as duplicated genes (17). Data on human ohnologs (gene duplicates retained after whole genome duplications) were collected from Makino et al., 2013 (64). The number of protein domains was derived from CDD (65). The gene chromatin state based on Hi-C experiments (19) was retrieved from the covariate matrix of MutSigCV v1.2.01 (2). Data on protein-protein and miRNA-gene interactions, gene evolutionary origin and gene expression were retrieved as described in An et al., 2016 (16). Briefly, human protein-protein interaction network was rebuilt from the integration of BioGRID v.3.4.125 (66); MIntAct v.190 (67); DIP (April 2015) (68); HPRD v.9 (69); the miRNA-gene interactions were derived from miRTarBase v.4.5 (70) and miRecords (April 2013) (60); gene evolutionary origin was assessed as described in D’Antonio et al., 2011 (18) using gene orthology from EggNOG v.4 (71); and gene expression in 30 normal tissues was retrieved from GTEx v.1.1.8 (72). Except gene length, duplication and ohnologs, all other systems-level properties had missing information for some of the 17,078 altered genes (Supplementary Table 1). To account for this, median imputation for continuous properties and mode imputation for categorical properties were implemented. Specifically, for each property median or mode values were calculated for known cancer genes and the rest of mutated genes. All missing values were replaced with their corresponding median or mode values.

### Application of sysSVM to EACs

The three steps of sysSVM were applied to 261 EACs (Figure 1A, Figure 1B, Supplementary Text). In step 1, all 34 features derived from molecular and systems-level properties (Supplementary Table 1) were mapped to the 17,078 altered genes in the cohort. Each feature was scaled to zero mean and unit variance to correct for the different numerical ranges across them. In step 2, 476 known cancer genes with damaging alterations (Supplementary Table 3) were used as a set of true positives for model selection. To optimise the parameters of the four kernels (linear, radial, sigmoid and polynomial) a grid search using 10,000 iterations of a three-fold cross validation was performed. At each iteration, the 476 known cancer genes were randomly split into 2/3 (around 317 genes) used as a training set and 1/3 (around 159 genes) used as the test set. At each increment of 100 cross validation iterations, the four best models (one per kernel) were chosen based on the median and variance of the sensitivity distribution across all previous iterations of cross-validation. The selection of the 100 sets of best models from all 10,000 cross-validation iterations was repeated 5 times, where all iterations were randomly re-ordered. In step 3, the resulting 500 best models were trained with the whole training set and used to rank the remaining 16,602 unique genes in each patient. A score was measured to combine the predictions from the four kernels and the genes not expressed in normal esophagus according GTEx annotation were excluded. These produced 500 lists of top 10 genes. Out of 500 best models, 38 had a unique set of parameters resulting in 24 unique lists of top 10 genes (Supplementary Table 4). These 24 lists ranged between 898 and 952 genes, with a core set of 598 genes shared across all of them. The most frequent top 10 list occurred 207 times (952_A, 41.4%, Supplementary Table 4). It was followed by 952_B (32.2%, 161 times) and 951_A (8.6%, 43 times). These three lists accounted for 82.2% of the 500 sets of top 10 genes, they shared 950 genes and were predicted by models differing in only one parameter (gamma in the polynomial kernel, Supplementary Table 4). Furthermore, the most frequent list was always predicted by the same set of best models. Therefore, 952_A represented a robust set of prediction and was considered as the final list of helper genes (Supplementary Table 5).

### Identification of perturbed processes and patient clustering

To identify the perturbed biological processes in the EAC cohort, both predicted cancer helper genes and known cancer driver genes were used. A manual revision of 476 known cancer genes altered in the ICGC cohort was performed and genes were considered as known drivers if (a) their somatic alteration had been previously associated with EAC, (b) they had a loss-of-function alteration and their tumour suppressor role had been reported in other cancer types (73), (c) they had a gain-of-function alteration and their oncogenic role had been reported in other cancer types (73). The resulting 202 known cancer drivers (Supplementary Table 3) and 952 cancer helpers were used for the gene set enrichment analysis against Reactome v.58 (74), composed of 1,877 pathways and 10,131 genes. After excluding pathways in levels 1 and 2 of Reactome hierarchy and those with less than 10 or more than 500 genes, 1,155 pathways were retained. These contained 9,061 genes, including 155 known drivers and 648 helpers. Gene set enrichment was assessed using a one-sided hypergeometric test and the resulting *P* values were corrected for multiple testing using the Benjamini & Hochberg method (Supplementary Table 6). Enriched pathways within the sets of known drivers or helpers were subsequently used to cluster samples taking into account the proportion of perturbed processes shared between samples. The Jaccard index (*A*) was calculated by deriving the proportion of shared perturbed processes between all possible sample pairs as:

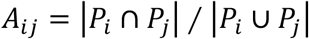

where *P_i_ and P_j_* are the perturbed processes in samples *i* and *j*, respectively.

Complete linkage hierarchical clustering using Euclidean distance between each row was performed on the resulting matrix. Clusters were visualised using ComplexHeatmap R package (75). To identify the optimal number of clusters, the median silhouette value of the samples for between 3 and 20 clusters was measured as a measure of clustering robustness (76).

### Analysis of RNA sequencing data

Purified total RNA was extracted from 92 EACs from the ICGC cohort and sequenced as described previously (10). RNA sequencing reads were then aligned to human reference genome hg19 and expression values were calculated using Gencode v19. The summariseOverlaps function in the R GenomicAlignments package was used to count any fragments overlapping with exons (parameters mode=Union, singleEnd, invertStrand and inter.feature were set according to the library protocol, fragments=TRUE, ignore.strand=FALSE). Gene length was derived as the number of base pairs in the exons after concatenating the exons per gene in non-overlapping regions. FPKM (Fragments Per Kilobase Million) were calculated for each gene as:

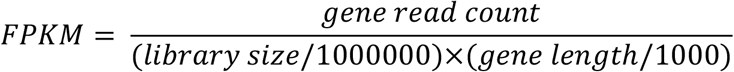

### Cell lines

Overexpression and editing experiments were carried out using the FLO-1 esophageal adenocarcinoma cell line obtained from the ECACC General Cell Collection. Cells were grown at 37°C and five per cent CO_2_ in DMEM + 2mM Glutamine + 10% FBS (Biosera) + 1/10,000 units of penicillin–streptomycin. Gene knockdown experiments were performed on OE19 cells obtained from the Francis Crick Institute cell services, OE33 cells obtained from the ECACC General Cell Collection and MFD1 cells obtained from the OCCAMS Consortium. OE19 and OE33 cells were grown in RPMI + 2mM Glutamine + 10% FBS (Biosera) + 1/10,000 units of penicillin–streptomycin. MFD1 cells were grown in DMEM + 2mM Glutamine + 10% FBS (Biosera) + 1/10,000 units of penicillin–streptomycin. All cells were maintained at 37°C and five per cent CO_2_, validated by short tandem repeat analysis and routinely checked for mycoplasma contamination.

### Gene overexpression

The vectors pCMVHA E2F1 (77) (Item ID 24225, Addgene), pLX_TRC317 (TRCN0000481188, Sigma-Aldrich) and pcDNA3.1+/C-(K)-DYK (Clone ID: OHu19407D, Genscript) were used to induce *E2F1*, *MCM7*, and *PAK1* overexpression, respectively. Cells were transfected according to the manufacturer’s protocol and selected with either G481/Geneticin (*E2F1*, *PAK1*) or Puromycin (*MCM7*). Empty vectors carrying G418 (pcDNA3.1+/C-(K)-DYK, Genscript) or Puromycin (Item ID 85966, Addgene) resistance were used as controls. The RNA from transfected cells was used to assess gene overexpression via quantitative RT-PCR using predesigned SYBR green primers (Sigma-Aldrich; Supplementary Table 7) and Brilliant III Ultra-Fast SYBR Green QRT-PCR Master Mix (Agilent Technologies). The average expression level across triplicates (*e*) was relativised to the average expression level of β-2-microglobulin (*c*):

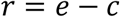

where *r* is the relative gene expression. The fold change (*fc*) between the relative gene expression after overexpression and the relative gene expression in the control condition (*r*_c_) was calculated as:

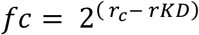

Each sample was assessed in triplicate and each experiment was repeated in biological duplicate.

### Gene editing

To induce *ABI2* and *NCOR2* gene knock-out (KO), the vector-free CRISPR-mediated editing approach was used as previously described (41). Briefly, cells were co-transfected using lipofectamine CRISPR max (Life technologies) with a 69-mer tracrRNA (Sigma-Aldrich), three gene-specific crRNAs (Sigma-Aldrich, Supplementary Table 7) and GeneArt Platinum Cas9 nuclease (Life technologies). To avoid off-target editing, all crRNAs used were verified to map only the gene of interest with a perfect match and additional hits in the genome with at least three mismatches. Control cells were transfected with the same protocol but using three non-targeting crRNAs. Gene editing was confirmed with Illumina Miseq sequencing. The regions surrounding the targeted sites were amplified from genomic DNA of edited cells with primers containing Illumina adapters (Supplementary Table 7) using Q5 Hot Start High-Fidelity 2X Master Mix (New England Biolabs). DNA barcodes were added with a PCR reaction before pooling the samples for sequencing on Illumina MiSeq with the 250 base-pair paired-end protocol. Sequencing reads were merged into single reads and aligned to the human reference genome hg19 using BBMerge and BBMap functions of BBTools (78), obtaining an average of 78,864 aligned reads per experiment. SNVs and small indels in the regions corresponding to each crRNA (Supplementary Table 7) were called using the CrispRVariants package in R (79) and the percentage of edited alleles was estimated as the percentage of variant reads in each experiment.

### Gene knockdown

Inducible gene knockdown was carried out using lentiviral pTRIPZ-TurboRFP (*MCM7*) or pSMART-TurboGFP (*PSMD3*) shRNA vectors (Dharmacon). For each gene, three shRNA vectors were tested (Supplementary Table 7). Virus was produced by co-transfecting HEK293T cells with pTRIPZ or pSMART constructs alongside psPAX2 and pMD2.G vectors (Addgene) using Fugene HD (Promega). Viral supernatant was collected at 24 and 48 hours and used for two rounds of infection of OE19, OE33 or MFD1 cells, using 8μg/ml hexadimethrine bromide (Sigma-Aldrich). Infected cells were selected after 48 hours with 2μg/ml puromycin for 7 days. To induce shRNA expression, cells were treated with 1μg/ml doxycycline (Sigma-Aldrich) for 16 hours. Gene expression with or without doxycycline was assessed by qRT-PCR using predesigned SYBR green primers (Sigma-Aldrich; Supplementary Table 7). Cells with the highest level of knockdown were then sorted by FACS to isolate medium expressing cells (the middle 30% of cells based on TurboRFP or TurboGFP fluorescence). Gene expression after sorting was measured by qRT-PCR 24 hours post-induction with 0-1μg/ml doxycycline, to determine the concentration of doxycycline required to reduce expression to levels equivalent to FLO-1 cells. The determined concentrations of doxycycline used for proliferation assays were 0.05μg/ml for OE19 PSMD3 shRNA3, 0.25μg/ml for OE33 PSMD3 shRNA3, 0.25μg/ml for MFD1 PSMD3 shRNA3, 0.75μg/ml for MFD1 MCM7 shRNA3.

### Cell proliferation

Cell proliferation was measured every 24 hours for three or four days, starting three hours after seeding the cells (time zero) using crystal violet staining, CellTiter 96 Non-Radioactive Cell Proliferation Assay (Promega) or CellTiter-Glo Luminescent Cell Viability Assay (Promega). Briefly, 4.5×10^3^ cells/well were seeded on 96-well plates in a final volume of 100μl per well. For inducible shRNA-expressing cells, doxycycline was added 48 hours prior to the start of the assay, and culture media replaced every 24-48 hours with fresh media containing doxycycline. For the CellTiter 96 Non-Radioactive Cell Proliferation Assay, 15 μl of the dye solution was added into each well and cells were incubated at 37°C for two hours. The converted dye was released from the cells using 100μl of the solubilisation/Stop solution and absorbance was measured at 570nm after one hour using the Paradigm detection platform (Beckman Coulter). For the CellTiter-Glo Luminescent Cell Viability Assay, 100μl of the CellTiter-Glo reagent was added into each well and luminescence was measured after 30 minutes using the Paradigm detection platform (Beckman Coulter). For all proliferation assays, four replicates per condition were measured at each time point and each measure was normalised to the average time zero measure for each condition. Each experiment was repeated at least twice. Conditions were compared using the two-tailed Student’s t-test.

### Flow cytometry

EdU incorporation and MCM loading were assessed using a modified version of the protocol described in Galanos et al., 2016 (80). Briefly, in each condition, 3×10^6^ cells were pulsed for 30 minutes with 10μM EdU (Invitrogen) before washing in 1% BSA/PBS. Chromatin fractionation was performed by incubating on ice for 10 minutes in CSK buffer (10mM HEPES, 100mM NaCl, 3mM MgCl_2_, 1mM EGTA, 300mM sucrose, 1% BSA, 0.2% Triton-X100, 1mM DTT, cOmplete EDTA-free protease inhibitor cocktail tablets, Roche). Cells were then fixed in 4% formaldehyde/PBS for 10 minutes at room temperature before washing in 1% BSA/PBS. Cells were permeabilised and barcoded (81) by incubating in 70% ethanol containing 0-15μg/ml Alexa Fluor 488 (Thermo Fisher) for 15 minutes, then washed twice in 1% BSA/PBS. Barcoded cells were subsequently pooled before incubating in primary antibody (mouse monoclonal anti-MCM7: Santa Cruz Biotechnology sc-56324, or rabbit polyclonal anti-MCM3: Bethyl Laboratories A300-192A) diluted 1:100 in 1% BSA/PBS for 1 hour. After washing in 1% BSA/PBS, samples were incubated for 30 minutes in secondary antibody (Alexa Fluor 555-conjugated donkey anti-mouse: A-31570, or donkey anti-rabbit: A-31572) diluted 1:500 in 1% BSA/PBS, then washed again in 1% BSA/PBS. EdU labelling with Alexa Fluor 647 azide was performed using Click-iT EdU flow cytometry assay kit (Invitrogen, C10424) following the manufacturer’s instructions before washing samples in 1% BSA/PBS. Samples were then incubated in 1% BSA/PBS containing RNase and 10mg/ml DAPI for 15 minutes before analysing with a BD LSR II Fortessa flow cytometer (BD Biosciences). Lasers and filters used include: 407nm laser with 450/50 bandpass filter; 488nm laser with 505 longpass and 530/30 bandpass filters; 561nm laser with 570 longpass and 590/30 bandpass filters; 640nm laser with 670/14 bandpass filter. Compensation was performed manually with single colour controls, using BD FACSDiva software (BD Biosciences). FlowJo 10.3 software was used to analyse MCM loading onto chromatin and EdU incorporation. Cells were gated to remove debris using FSC-A/SSC-A, then gated to isolate singlets using DAPI-H/DAPI-A (Supplementary Figure 9). The cells were then separated by gating the barcoded populations using 488-A/DAPI-A. Cells were finally separated into cell cycle gates (G1, S1-4, G2) based on EdU-647-A and DAPI-A (SupplementaryFigure 9), and the geometric mean fluorescence intensity was obtained for each channel (MCM-555 or EdU-647).

## AUTHOR CONTRIBUTIONS

FDC conceived and directed the study. AM developed all the bioinformatics methods under the supervision of FDC, MC and CY. LB and EF performed the experiments with the help of MH and PS. AM, LB, EF, FDC analysed the data. JP helped with the analysis of RNA-Seq data. FDC, AM, LB and EF wrote the manuscript. RCF, PS, MH, JL edited the manuscript. All authors approved the manuscript.

## ACKNOWLEDGMENTS

We thank Stephanie Hills and John Diffley (The Francis Crick Institute) for their help with the experiments on E2F1 and MCM7 and for the discussion of results. We thank Valdone Maciulyte (The Francis Crick Institute) for help with the MiSeq library preparation. We are grateful for the support of the High Throughput Screening Facility, Flow Cytometry Facility, Advanced Sequencing Facility and Cell Services of the Francis Crick Institute.

## LIST OF SUPPLEMENTARY DATA

**Supplementary Text.** Description of the sysSVM algorithm

**Supplementary Figure 1.** Pairwise correlation between 34 features for classification

**Supplementary Figure 2.** tSNE plots of 34 properties of known cancer genes

**Supplementary Figure 3.** sysSVM validation

**Supplementary Figure 4.** Co-amplifications and features of known cancer drivers

**Supplementary Figure 5.** EAC clustering using pathways enriched in known drivers

**Supplementary Figure 6.** Identification of the optimal number of clusters

**Supplementary Figure 7.** EAC clustering using pathways enriched in helpers

**Supplementary Figure 8.** Comparison of helpers using different ranking cut offs

**Supplementary Figure 9.** Flow cytometry gating strategy and pseudocolour plots

**Supplementary Table 1.** Description of sysSVM features

**Supplementary Table 2.** Description of EACs used in this study

**Supplementary Table 3.** Known cancer genes with driver alterations

**Supplementary Table 4.** Selection of best models and final list of helper genes

**Supplementary Table 5.** Cancer helper genes in 261 EACs

**Supplementary Table 6.** Gene set enrichment analysis of EAC drivers or helpers

**Supplementary Table 7.** List of oligos used in the study

